# Temperature-Related Intensity Change (TRIC)-based High Throughput Screening Platform for the Discovery of CHI3L1-Targeted Small Molecules

**DOI:** 10.1101/2025.08.13.670152

**Authors:** Longfei Zhang, Moustafa T. Gabr

## Abstract

Chitinase-3-like 1 (CHI3L1) protein is a secreted glycoprotein involved in various normal physiological processes, while its abnormal elevation is closely associated to carcinogenesis. CHI3L1 promotes tumor progression by recruiting and polarizing immune cells to sustain an immunosuppressive microenvironment, while directly stimulating cancer cell proliferation and migration. Although CHI3L1 deletion has shown therapeutic potential in animal models, the development of small molecule modulators remains limited. To address this gap, we developed a TRIC-based high-throughput screening (HTS) platform for CHI3L1 binding and applied it to a 5,280-compound library, yielding 11 binders (hit rate: 0.21%). Of these, three compounds (**9N05, 11C19**, and **3C13**) were validated by surface plasmon resonance (SPR). Among the validated hits, **9N05** showed the strongest binding affinity to CHI3L1 (K_d_ = 202.3 ± 76.6 μM) but only weak inhibition of CHI3L1/galectin-3 interaction. In contrast, **11C19** significantly disrupted this interaction (73.9 ± 7.9% inhibition at 100 μM; IC_50_ = 188.6 ± 0.18 μM). This proof-of-concept study establishes TRIC-based screening as a viable approach for identifying CHI3L1-targeted small molecules and supports its use in future modulator discovery.

## INTRODUCTION

Chitinase-3-like 1 (CHI3L1, also known as YKL-40), a member of the glycoside hydrolase family 18, is a secreted glycoprotein that plays important roles in apoptosis, cell migration, inflammation modulation, and angiogenesis.^1-4^ Due to a mutation in its catalytic domain, CHI3L1 lacks the enzymatic activity characteristic of other family members, but retains the ability to interact with multiple receptor proteins—such as IL-13Rα2, galectin-3, and TMEM219—and functions as a signaling molecule.^5-8^ These interactions activate downstream pathways including ERK1/2, AKT, WNT/β-catenin, and NF-κB, thereby regulating diverse physiological processes and modulating the immune microenvironment.^3, 9, 10^

In recent decades, CHI3L1 has emerged as both a diagnostic biomarker and a potential therapeutic target in cancer, as accumulating evidence has revealed its critical role in tumorigenesis. According to clinical studies, abnormally elevated CHI3L1 levels can be detected in the serum of patients with various types of cancer, including non-small cell lung cancer, hepatocellular carcinoma, and glioblastoma.^6, 11-13^ Moreover, serum CHI3L1 levels demonstrated a positive correlation with disease severity, and high CHI3L1 expression is generally associated with poor prognosis and reduced survival.^14-17^ Besides, elevated CHI3L1 expression was observed in neurodegenerative diseases, such as Alzheimer’s disease, and autoimmune disorders, including rheumatoid arthritis.^6, 18^

During carcinogenesis, CHI3L1 recruits immune cells—such as macrophages and neutrophils— into the tumor microenvironment and stimulates them to release pro-inflammatory cytokines, including IL-6, IL-8, and TGF-β, thereby establishing an inflammatory environment that supports cancer cell proliferation and migration.^19, 20^ CHI3L1 also attracts tumor-associated macrophages (TAMs) and polarizes them into an M2-like phenotype via an IL-13Rα2-dependent signaling pathway, contributing to an immunosuppressive microenvironment that dampens antitumor immunity and facilitates tumor immune evasion.^14, 21^ Additionally, CHI3L1 impairs immune responses by skewing the Th1/Th2 balance toward Th2 dominance through IFN-γ- related signaling.^19^ In 2018, Kim et al. intravenously injected B16F10 melanoma cells into CD4- specific CHI3L1-knockout mice and observed a reduced number of lung metastases, accompanied by decreased vascular infiltration and elevated expression of IFN-γ and TNF-α compared to controls.^22^ Subsequent qPCR analysis revealed increased expression of Th1-related mRNAs in the tumor region, indicating a tumor-suppressive immune environment.^22^ Moreover, CHI3L1 has been shown to directly promote cancer cell proliferation and metastasis. In another 2018 study, Qiu et al. silenced CHI3L1 in HepG2 and Bel-7404 hepatocellular carcinoma cells, resulting in significantly reduced migration *in vitro*.^23^ Following intravenous administration, mice injected with CHI3L1-knockdown cells developed fewer distant metastases compared to controls (3 – 4 vs. 6 lesions), supporting the critical role of CHI3L1 in tumor progression and highlighting its potential as a therapeutic target in cancer treatment.^23^

In 2024, Czestkowski et al. reported two small-molecule inhibitors, compounds **30** and **36**, obtained from an AlphaScreen-based high-throughput screening (HTS) assay targeting the interaction between CHI3L1 and a fractionated heparan sulfate polymer (HSIII).^24^ Both compounds demonstrated potent inhibitory activity, with IC_50_ values of 37 nM and 26 nM, respectively.^24^ Notably, compound **36** exhibited a maximum inhibition rate of 83%, compared to 27% for compound **30. K284-6111** was discovered by Prof. Hong’s laboratory through a structure-based virtual screening approach, which displayed efficient inhibition ability on the growth of A549 and H460 cells via disturbing the interaction between CHI3L1 and IL-13Rα2.^25^

In the following *in vivo* study. In subsequent *in vivo* experiments, intravenous administration of **K284-6111** (0.5 mg/kg, 3-day intervals for 3 weeks) led to a reduction in metastatic number in tumor-bearing mice (B16F10 and A549 models), compared to the vehicle-treated group.^25, 26^ Interestingly, oral administration of **K284-6111** (3.0 mg/kg for 4 weeks) in Tg2576 mice alleviated memory impairment symptoms according to the water maze test, and ELISA analysis further confirmed a reduction in Aβ deposition.^27^

Even though CHI3L1 inhibition demonstrated promising therapeutic effects in animal studies for cancer treatment, as the research is still in a relatively early stage, inhibitors used in most studies are limited to antibodies (mAbs), with only a few molecular inhibitors or binders for CHI3L1 reported, and none have been approved for clinical trials. Compared to mAbs, molecular medicine demonstrates advantages, including improved tissue penetration, friendly administration, oral bioavailability, and low manufacturing costs. Besides, owing to the facility in structure modification, long-term side effects could be reduced via binding-specificity and pharmacokinetics optimization. To address the lack of molecular CHI3L1 inhibitors, developing novel platforms for HTS is necessary and urgent. Temperature-related intensity change (TRIC) is a fluorescence-based technology designed for protein interaction detection. Compared with commonly used HTS platforms, such as TR-FRET, SPR, and AlphaLISA, TRIC demonstrated distinct advantages in efficiency, sensitivity, and economy. For each sample, only picomolar protein is required, and a 384-well plate could be measured within 40 min with dual laser cooperation, making it an ideal platform for HTS of proteins, especially for large-scale screening studies.

In this study, using the TRIC technology, 5280 small molecules from the Discovery Diversity Set (DDS-10, Enamine, Kyiv, Ukraine) were screened for CHI3L1 binding (**Figure 1a**), followed by the control study and binding validation study on the same platform, 11 molecules were identified as hits, yielding a hit rate of 0.21%. Following orthogonal validation studies were performed on microscale thermophoresis (MST) and surface plasmon resonance (SPR), and three compounds were verified as binders to CHI3L1 with moderate binding affinities (K_d_ = 202.3 ± 76.6 μM, 480.7 ± 403.9 μM, and 202.3 ± 76.6 μM for **9N05, 11C19**, and **3C13**, respectively). Moreover, in the AlphaLISA-based inhibition assay, **11C19** demonstrated inhibitory activity against the CHI3L1/galectin-3 interaction, with a maximum inhibition rate of 73.9 ± 7.9%.

**Figure 1.**
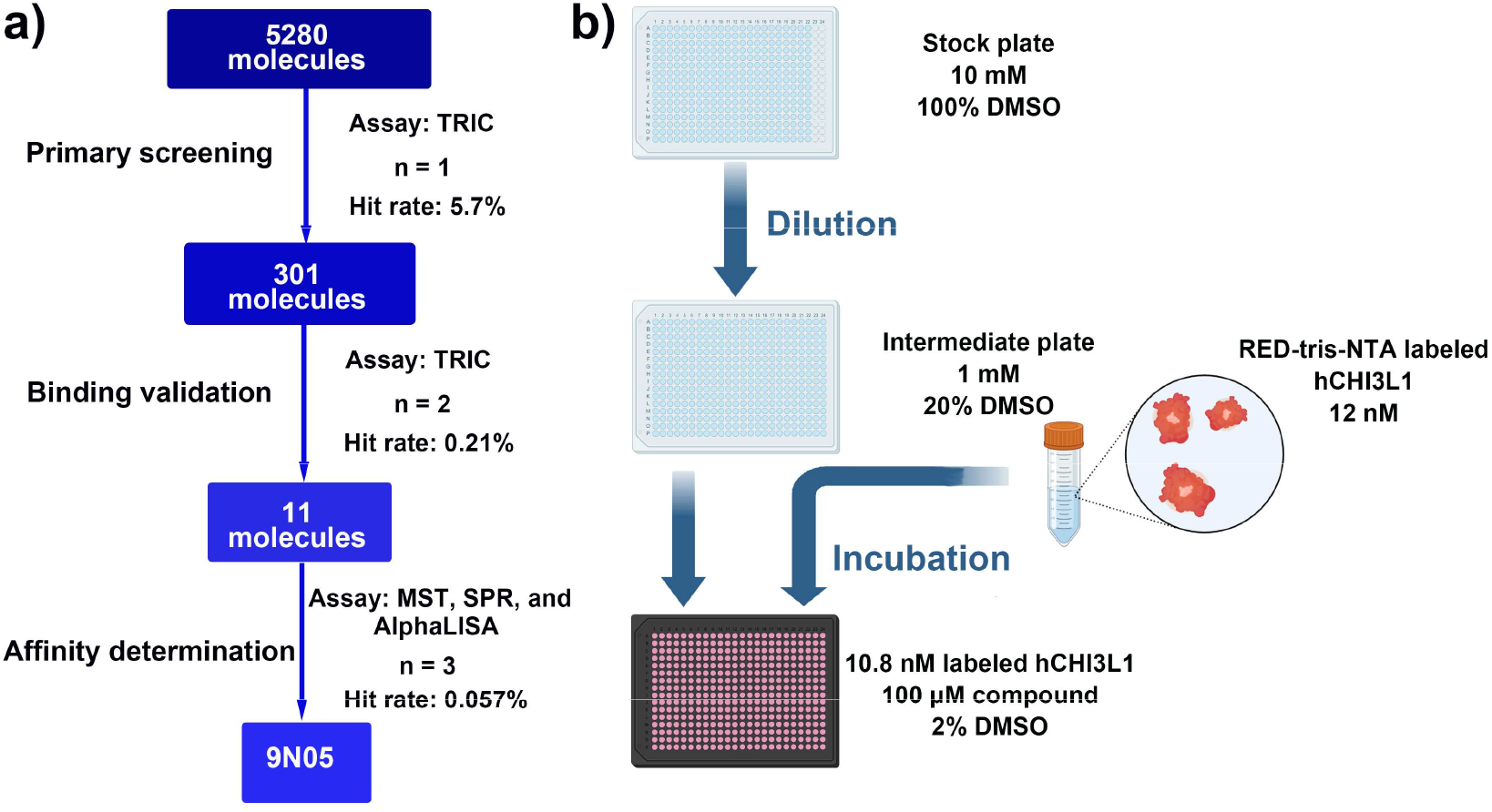
Workflow for the discovery of CHI3L1-targeted small molecules. **a)** Schematic representation of HTS workflow. **b)** Schematic representation of workflow for the TRIC-based assay.

## METHODS

### TRIC-based HTS

The HTS was performed using a TRIC-based method on a Dianthus NT.23 Pico (NanoTemper Technologies, Germany), and 5280 molecules from the DDS-10 library were screened.

hCHI3L1-His protein (CH1-H5228, Acro Biosystems, Newark, DE, USA) was labeled with the Monolith His-Tag Labeling Kit RED-tris-NTA 2nd Generation (MO-L018, NanoTemper Technologies, München, Germany) following the manufacturer’s instructions. Briefly, 100 nM dye was mixed with 200 nM hCHI3L1 in a 1:1 volume ratio, and incubated at room temperature (r.t.) for 30 min in the dark before each experiment.

As shown in **Figure 1b**, 1 μL of compound DMSO solution (10 mM) was transferred from the stock plate to an intermediate 384-well plate with a MINI 96 channel portable electronic pipette (Integra Bioscience, Zizers, Switzerland), and 1 μL DMSO and 8 μL assay buffer were added subsequently. 3 μL of the above solution was mixed with the freshly prepared labeled hCHI3L1 with a final concentration of 100 μM and 10.8 nM for candidate molecule and protein, respectively. The mixture was incubated at r.t. for 30 min in darkness, and centrifuged at 1000 x g before loading into the Dianthus NT.23 Pico to measure the normalized fluorescence (F_norm_). Assay buffer: 10 mM HEPES, 150 mM NaCl, 1% Pluronic F127, 1 mM TCEP, 2.5% DMSO, pH = 7.4. Assay buffer with DMSO was used as the negative control.

Data was analyzed on the DI. Screening Analysis software (NanoTemper Technologies, Germany).

### Control Study

1 μL of compound DMSO solution (10 mM) was transferred from the stock plate to an intermediate 384-well plate and mixed with 1 μL DMSO and 8 μL assay buffer. 1 μL of the above solution was mixed with the RED-tris-NTA 2nd Generation dye, with a final concentration for compound and dye of 100 μM and 5.4 nM, respectively. The mixture was incubated at r.t. for 30 min in darkness, and centrifuged at 1000 x g before loading into the Dianthus NT.23 Pico to measure the normalized fluorescence (F_norm_). Assay buffer: 10 mM HEPES, 150 mM NaCl, 1% Pluronic F127, 1 mM TCEP, 2.5% DMSO, pH = 7.4. The assay was performed in technical duplicate, and the data were analyzed on the DI. Screening Analysis software.

### TRIC-based Binding Validation

The same method in TRIC-based HTS was used.

### MST-based Affinity Measurement

The protein was labeled with the RED-tris-NTA 2nd Generation dye as described above. Gradient concentrations of compounds were prepared in the assay buffer and incubated with labeled protein (final concentration: 20 nM) at r.t. for 30 min in dark. The solution was loaded into a Monolith Premium capillary (MO-K025, NanoTemper Technologies, Germany) and measured on a Monolith NT.115 instrument (NanoTemper Technologies, Germany).

### SPR-based Affinity Measurement

The assay was conducted on a Biacore™ 8K SPR system (Cytiva, Marlborough, MA, USA) at 25 °C. hCHI3L1-His protein (8.0 μg/mL in acetate buffer, pH 5.0) was labeled on a Series S Sensor Chip CM5 (29104988, Cytiva, Marlborough, MA, USA) using a commercial amine coupling kit (BR100050, Cytiva, Marlborough, MA, USA). Immobilization was performed at a flow rate of 10 μL/min for 420 s, reaching an immobilization level of approximately 3000 response units (RU), followed by blocking with ethanolamine. An ethanolamine-blocked flow cell on the same sensor channel was used as a reference. Immobilization buffer: PBS-P+ (28995084, Cytiva, Marlborough, MA, USA)

Gradient concentrations of the tested compound were prepared in the assay buffer supplemented with 5.0% DMSO and injected over the sensor chip in a single-kinetic cycle model, using a flow rate of 30 μL/min for 120 s per injection. After each injection, a dissociation phase of 600 seconds was performed, followed by chip regeneration using regeneration buffer (10 mM sodium acetate, 250 mM NaCl, pH 4.0) with a flow rate of 30 μL/min for 30 s.

Data were analyzed using Biacore™ Insight Evaluation Software (Cytiva, Marlborough, MA, USA). The assay was performed in individual triplicate and the K_d_ values were given as mean ± SD.

### AlphaLISA-based Assay

Single-dosage (final concentration: 100 μM) or gradient concentrations of tested compounds were prepared in the assay buffer (AL018C, Revvity, Inc., Waltham, M.A., U.S.) supplemented with 2.5% DMSO and incubated with CHI3L1-His (CH1-H5228, Acro Biosystems, Newark, DE, USA) and galectin-3-GST (Cat # 10289-H09E, Sino Biological, Beijing, China), and the final concentration of two proteins were both 60 nM. The mixture was incubated at r.t. for 3 h in dark. AlphaScreen anti-6His donor beads (15 μg/mL, AS116D, Revvity, Inc., Waltham, M.A., U.S) and AlphaLISA anti-GST acceptor beads (15 μg/mL, AL110C, Revvity, Inc., Waltham, M.A., U.S) were added, followed by an extra hour incubation at r.t. in darkness. The signal detection was performed using a Tecan Spark. plate reader (Tecan Group Ltd., Männedorf, Switzerland) using the AlphaLISA model (wavelength: 623 nm; bandwidth: 25 nm). Assay buffer (2.5% DMSO) with CHI3L1-His and galectin-3-GST was performed as the positive control, while buffer with CHI3L1-His solely as the negative control. The inhibition rate was calculated as: inhibition rate = (fluorescence_sample_ - fluorescence_positive_) / (fluorescence_negative_ - fluorescence_positive_) × 100%. The assay was performed in technical triplicate and results were given as mean ± SD.

## RESULTS AND DISCUSSION

### TRIC-based Primary Screening

Aiming to discover novel small molecule binders of CHI3L1, a TRIC-based HTS was performed on 5280 molecules from the DDS-10 (**Figure 1a**). To minimize reagent consumption and cost, the primary screening was performed in singlicate at a single dosage (100 μM). After incubating with fluorophore-labeled CHI3L1 (10.8 nM) for 30 min in the dark, the normalized fluorescence intensity (F_norm_) was measured to evaluate the interaction with protein (**Figure 1b**), and assay buffer with 2% DMSO served as the negative control. Due to the slight difference in power efficiency between the two lasers in the Dianthus instrument, F_norm_ values were only compared to negative controls measured using the same laser.

In the primary screening, to account for potential signal variation caused by the singlicate measurements, a relatively permissive threshold was applied, and all molecules with the F_norm_ values falling outside of the range of 5 times the standard deviation (SD) from the corresponding negative control were treated as potential hits and selected for the following studies. As shown in **Figure 2**, 577 compounds displayed an obvious signal shift compared to the corresponding negative control, and the hit rate was calculated as 10.9% in this step.

**Figure 2.**
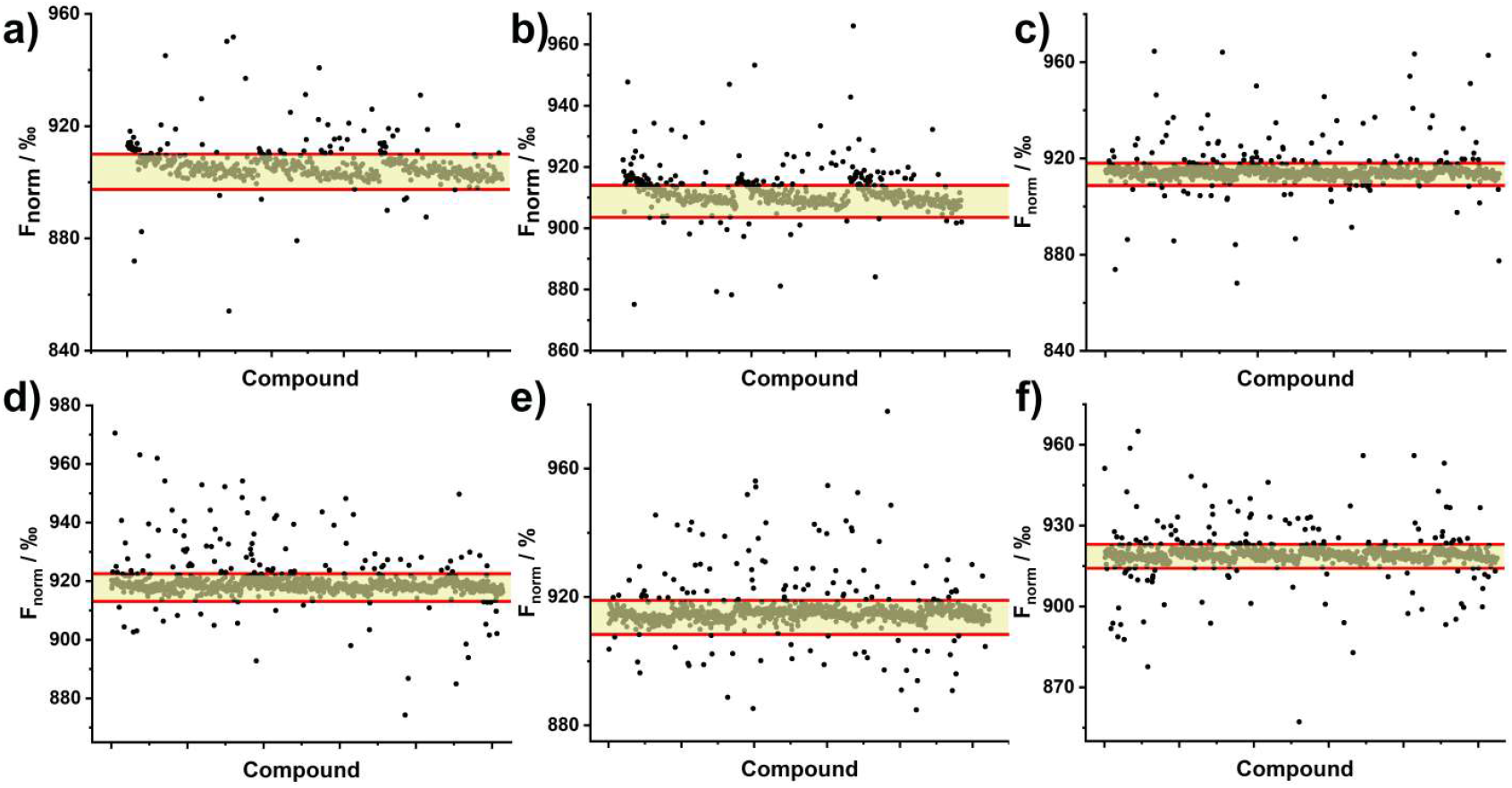
Primary screening results from the HTS campaign for the discovery of small molecule CHI3L1 binders using TRIC. Results of the primary screening at 100 μM. Pale yellow region between the red lines indicates the range within 5 × SD from the mean F_norm_ value of negative controls. The graphs show the results from a single experiment.

However, since the TRIC is a fluorescence-based technology, molecules with fluorescence or label interaction may interfere with the measurement, resulting in false-positive results. For filtering out false-positive candidates, the initial fluorescence intensities of tested compounds were examined. A range of 3 times SD from the mean value of the negative control was set as the threshold, and molecules with initial fluorescence intensities outside of this range were flagged as suspect. Among the 577 hits identified from the primary screening, 352 candidates displayed shifted fluorescence intensities from negative controls, which were subjected to a subsequent control study to assess the signal reliability. After incubating with the fluorophore (5.4 nM) alone under the same conditions as the primary screen, only 76 compounds displayed close F_norm_ values and fluorescence intensities to the negative control, while the remaining showed significant shift in at least one of these parameters (**Figure S1, Supporting Information, SI**).

Based on the primary screening and a follow-up control study, 301 candidates were selected kept for the following screening, and the hit rate was 5.7%.

### TRIC-based Binding Validation

To validate the CHI3L1 binding affinity of the 301 compounds identified from the previous step, a TRIC-based binding validation was performed at a single dosage (100 μM, n = 2) under the same conditions as the primary screening, and a similar control study was performed as described previously. To elevate the reliability of identified hits, a threshold of ±10 SD from the mean F_norm_ value was applied, and molecules with initial fluorescence intensities above 5 times SD from the negative control were excluded.

As shown in **Figure 3**, most compounds from the previous steps were filtered out as the weak F_norm_ shift (spots in the pale yellow area) or obvious deviation in initial fluorescence after incubation (spots in orange) under the strict criteria, and only 11 candidates (spots in blue) were identified as hits (hit rate: 0.21%).

**Figure 3.**
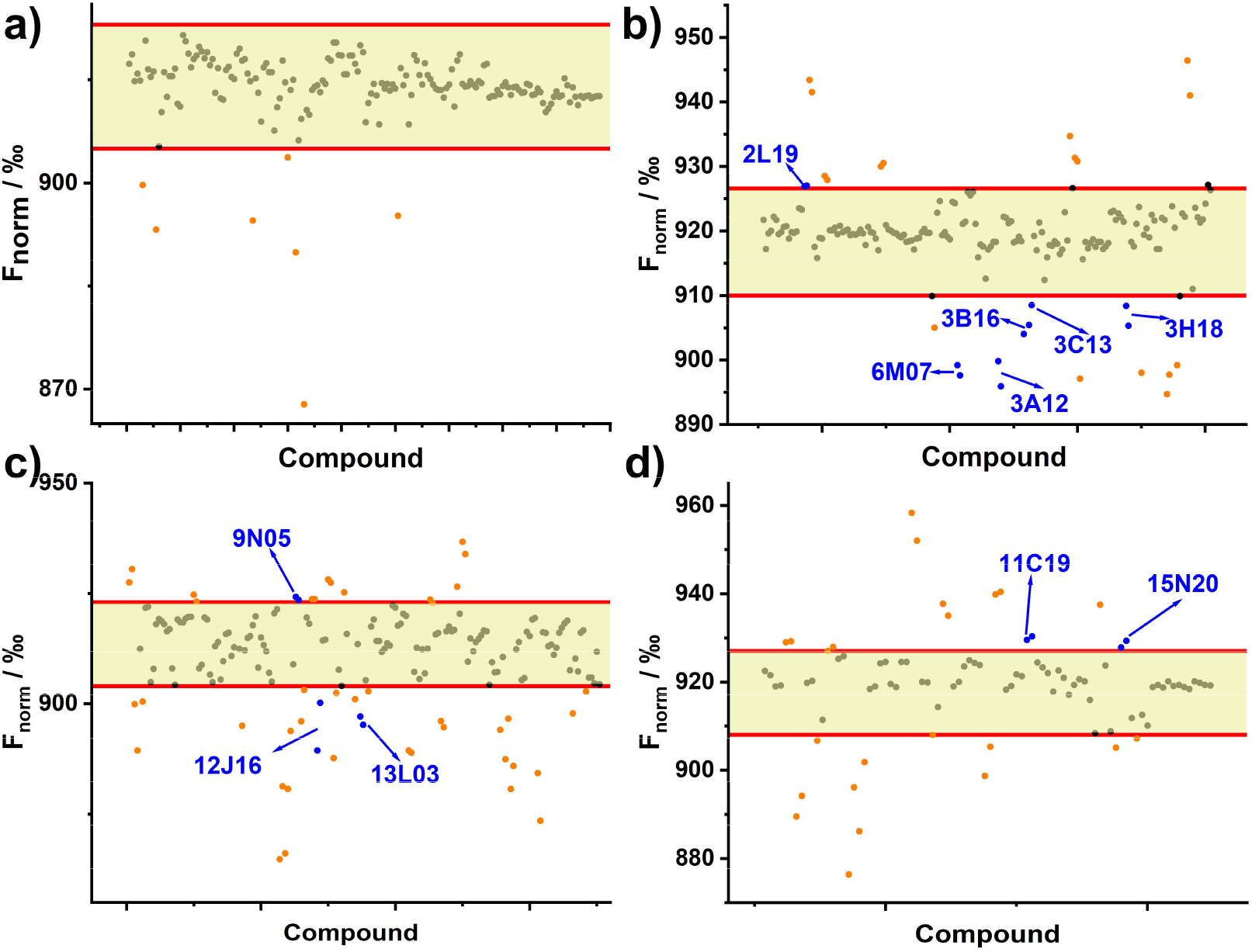
Binding validation of small molecule CHI3L1 hits identified from the primary TRIC screening. Results of the binding validation at 100 μM. Pale yellow region between the red lines indicates the range within 10 × SD from the mean F_norm_ value of negative controls. Black spot indicates compounds fell inside of the negative range, orange spots indicate compounds showed signal shift in control study, and blue spots indicate hit identified. F_norm_ was measured in duplicate and both individual replicate values are plotted.

### Biological Properties Assessment

The binding affinities of the validated 11 hit compounds (**Table 1**) were then evaluated using MST. After incubating with fluorophore-labeled CHI3L1 (20 nM) for 30 min in the dark, gradient concentrations of tested compounds were loaded into the Monolith Premium capillary, and the F_norm_ were recorded. Even though all 11 compounds demonstrated positive signals in previous screening studies, only five compounds (**9N05, 11C19, 2L19, 12J16**, and **15L20**) demonstrated clear dosage-dependent signal changes, while signals of other candidates showed high variability without a defined trend with increasing concentrations (**Figure 4c, 4f, 4i**, and **S3, SI**). However, owing to the limited aqueous solubility, the maximum concentrations of tested molecules could only reach 250 μM, which prevented signal saturation within the measurement range, thus only approximate K_d_ value could be determined for compound **9N05, 11C19**, and **15L20** (61.4 μM, 125.0 μM, and 139.0 μM, respectively), while **3C13, 2L19**, and **12J16** failed to give any values (**Figure 4c, 4f, 4i**, and **S3, SI**).

**Table 1.**
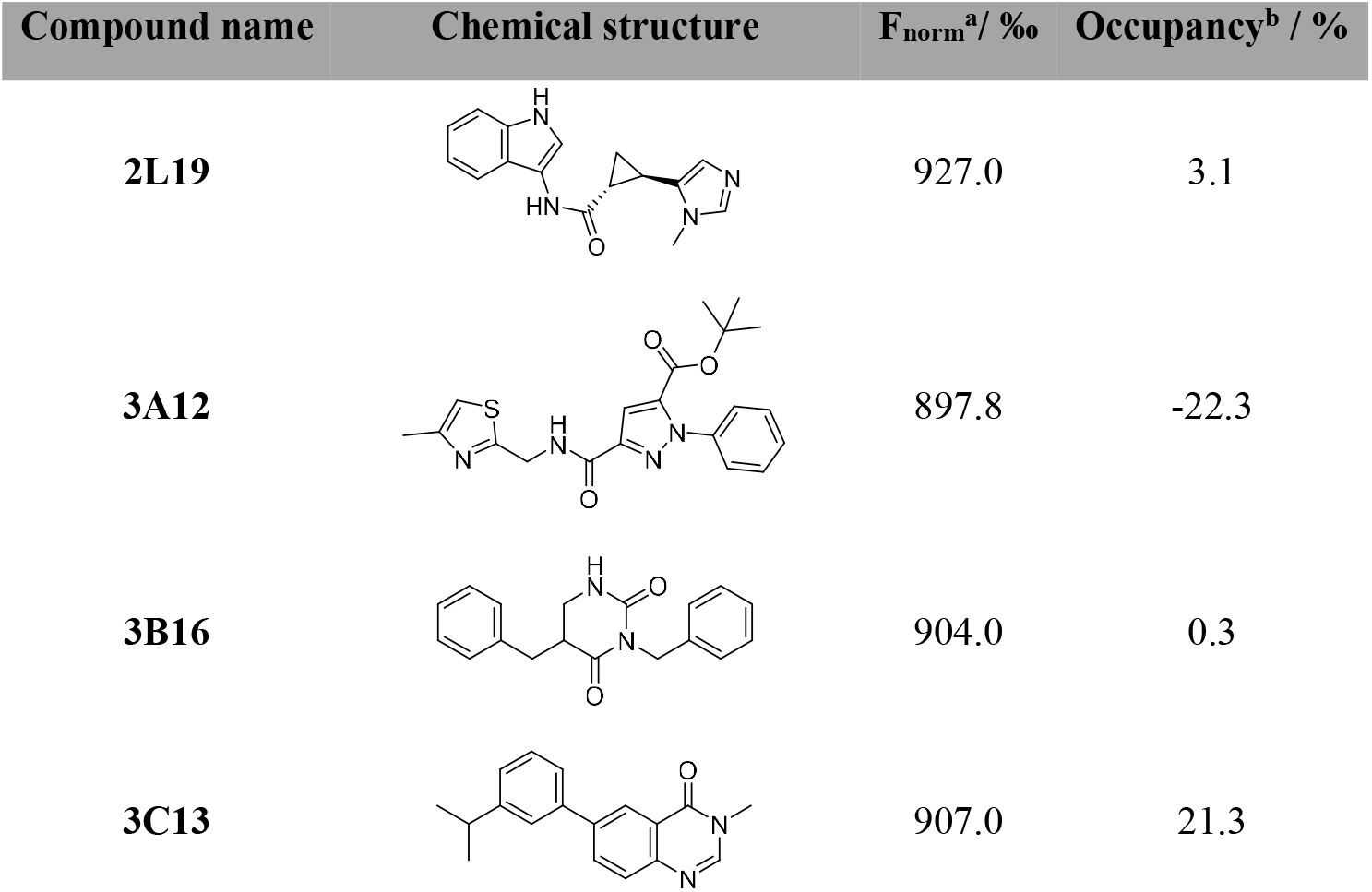
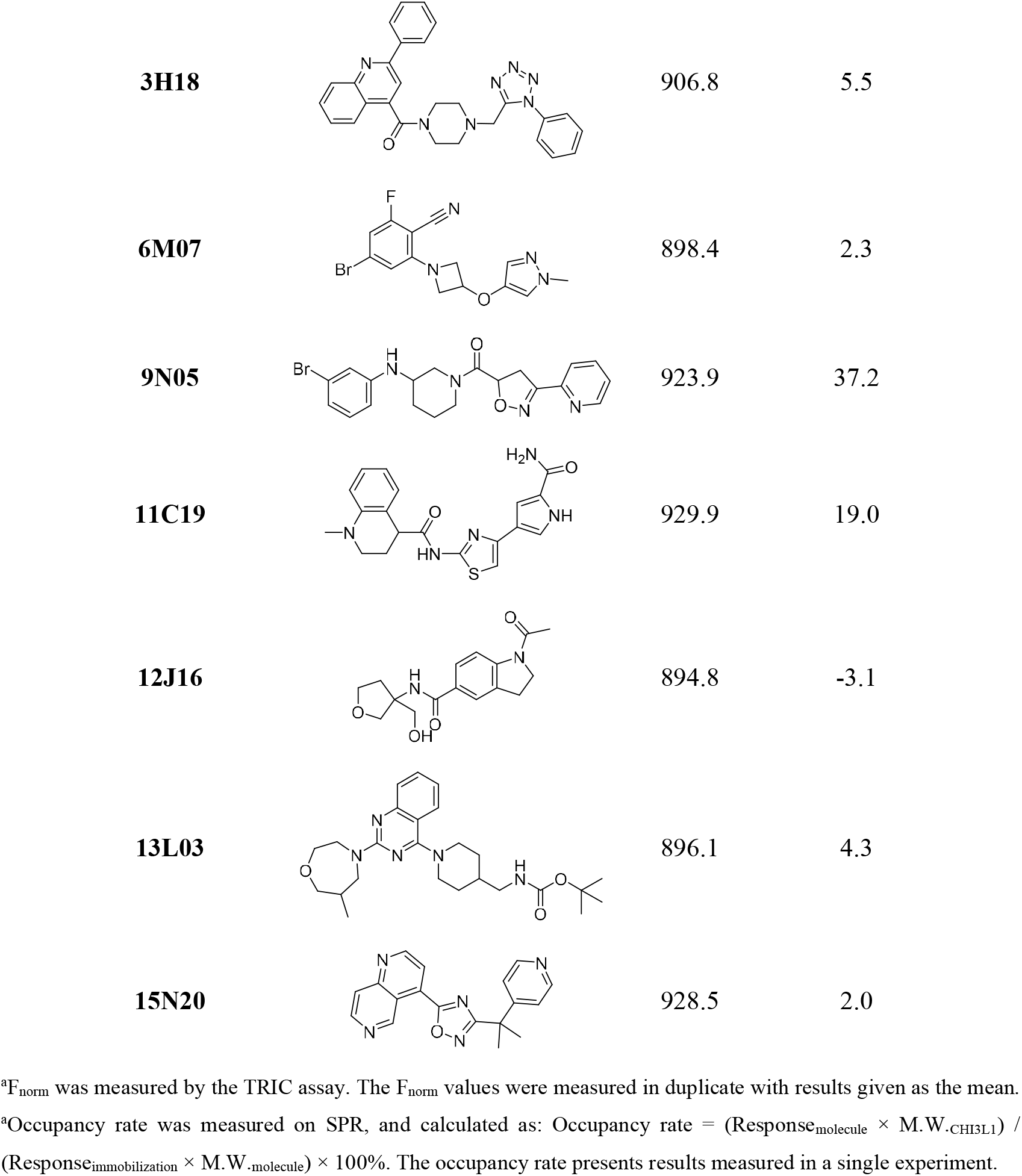
Chemical structures, F_norm_ values and occupancy rates of 11 CHI3L1 hits identified in the TRIC-based assay.

**Figure 4.**
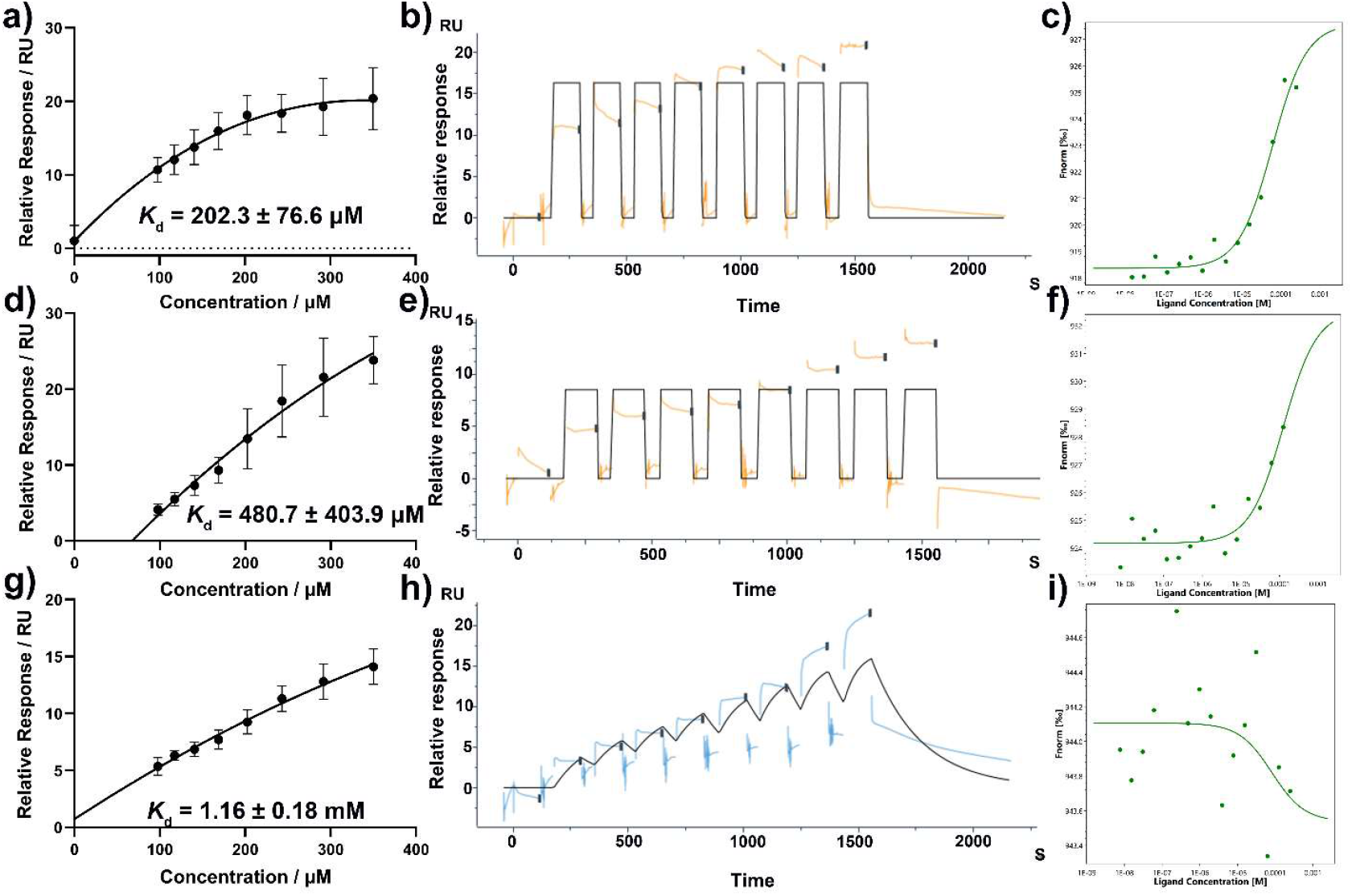
Orthogonal validation of CHI3L1 hits using SPR. Orthogonal binding validation of **9N05** (**a** - **c**), **11C19** (**d** - **f**), and **3C13** (**g** - **i**) using SPR (**a, b, d, e, g**, and **h**) and MST (**c, f**, and **i**). The relative responses in SPR (**a, d**, and **g**) were measured in triplicate with results given as the mean ± SD. The sensorgram graphics (**b, e**, and **h**) represent results of individual triplicate experiments. F_norm_ in MST was measured in a single experiment.

To orthogonally validate the CHI3L1 binding affinity of the 11 hit compounds, the compound- CHI3L1 interaction was assessed using SPR. After immobilizing CHI3L1 on a Series S CM5 Chip with an immobilization level at approximate 3000 RU, 100 μM compound (n = 1) was injected over the chip at a rate of 30 μL/min for 420 s, and the occupancy rates (occupancy rate = (Response_molecule_ × M.W._CHI3L1_) / (Response_immobilization_ × M.W._molecule_) × 100%) was recorded to assess the binding (**Table 1**). Among 11 hits, 3 compounds were confirmed as binders of CHI3L1, and the occupancy rates were calculated as 37.2%, 19.0%, and 21.3% for **9N05, 11C19**, and **3C13**, respectively. In the following dosage-response assay, **9N05** and **11C19** demonstrated a fast association followed by a quick dissociation process, and clear saturation plateaus were obtained at every concentration (**Figure 4b** and **4e**). After a 600-s wash with running buffer, the sensorgram returned to baseline, indicating the specific binding of both compounds. In contrast, **3C13** reached equilibrium rapidly at concentrations below 178.5 μM, while the signal continued to rise until the end of injection at 250 μM and 350 μM (**Figure 4h**). The response signal demonstrated a slow decline in the following wash process, and remained at about 3 RU until the end, indicating that non-specific binding happened at high concentrations (250 μM and 350 μM). The *K*_d_ values of **9N05, 11C19**, and **3C13** were calculated as 202.3 ± 76.6 μM, 480.7 ± 403.9 μM, and 1.16 ± 0.18 μM, respectively (**Figure 4a, 4d**, and **4g, Table 1**).

Galectin-3 is a β-galactoside-binding protein that interacts with CHI3L1 and plays an important role in the pathological processes of carcinogenesis, such as immune suppression, M2 macrophage polarization, and angiogenesis. In this study, an AlphaLISA assay was established to evaluate the inhibitory activities of **9N05, 11C19**, and **3C13** against the CHI3L1/galectin-3 interaction.

After incubating 100 μM compound (n = 3) with CHI3L1-His, galectin-3-GST, anti-His-donor beads, and anti-GST-acceptor beads in the dark, the inhibition rate was measured as described previously. Interestingly, although compound **9N05** demonstrated more potent binding affinity than other two candidates in both MST and SPR assays, only weak inhibitory effect was detected against the CHI3L1/galectin-3 interaction at 100 μM (inhibition rate: 21.5 ± 2.6%), while **11C19** displayed an unexpectedly potent inhibition ability at the same concentration, with an inhibition rate of 73.9 ± 7.9% (**Figure 5a**). Compound **3C13** also displayed a weak inhibitory activity, and the inhibition rate was calculated as 27.5 ± 3.1%. In the following dosage-response assay (n = 3), **11C19** demonstrated a clear dosage-dependent increase in inhibition with increasing concentrations, and the IC_50_ value was calculated as 188.6 ± 97.8 μM (**Figure 5c**), while only scattered signals could be found for the other two compounds (**Figure 5b** and **5d**).

**Figure 5.**
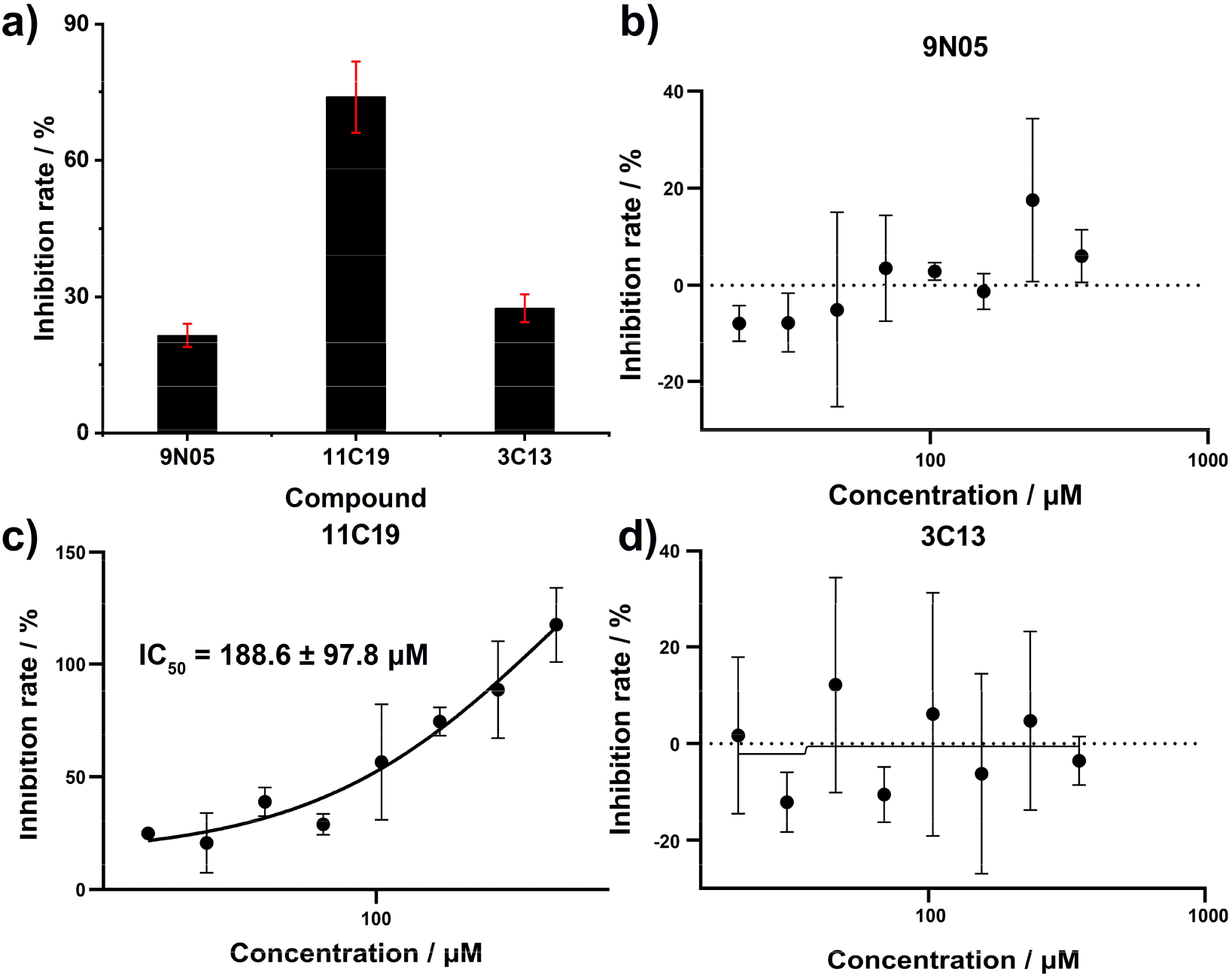
AlphaLISA-based inhibition assay of CHI3L1/galectin-3 interaction by CHI3L1 hits. **(a)** Single dosage (100 μM) inhibitory activity check of **9N05, 11C19**, and **3C13. (b - d)** Inhibition curve of **9N05** (**b**), **11C19** (**c**), and **3C13** (**d**) against CHI3L1/galectin-3 interaction using the ApphaLIS inhibition assay. The inhibition rates were measured in triplicate with results given as the mean ± SD.

In summary, a TRIC-based HTS platform was developed and employed to screen a small molecule library containing 5280 candidates for the ability to bind CHI3L1. The screening platform demonstrated intrinsic advantages in efficiency, economy, and sensitivity. Followed by a control study and binding validation on the same platform, 11 hits with novel and diverse chemical scaffolds were identified, corresponding to a hit rate of 0.21%. Based on the following orthogonal validation study using SPR, **9N05, 11C19**, and **15L20** were confirmed as CHI3L1 binders with *K*_d_ values of 202.3 ± 76.6 μM, 480.7 ± 403.9 μM, and 1.16 ± 0.18 mM, respectively, and the final hit rate of this screen is 0.057%. The inhibitory activities of three compounds against CHI3L1/galectin-3 interaction were evaluated using an AlphaLISA assay, in which the protein interaction was strongly disturbed by **11C19** (inhibition rate: 73.9 ± 7.9%), with an IC_50_ of 188.6 ± 0.18 μM. Collectively, this work validated the feasibility and reliability of the TRIC-based HTS platform for CHI3L1-binder screening, and provides a foundation for future development of CHI3L1-targeted therapeutics.

## AUTHOR INFORMATION

## Corresponding Author

Moustafa T. Gabr

## CRediT Authorship Contribution Statement

Longfei Zhang: Conceptualization, Methodology, Validation, Formal Analysis, Investigation, Data Curation, Writing – Original Draft, Visualization. Moustafa T. Gabr: Writing - Review & Editing, Funding Acquisition.

## Data Availability Statement

The data that support the findings of this study are available in the Supporting Information.

## ACKNOWLEDGMENT

We gratefully acknowledge financial support from the National Institute of Neurological Disorders and Stroke (Award ID: R01NS136524).

## ABBREVIATIONS

CHI3L1: Chitinase-3-like 1;
DMSO: dimethyl sulfoxide;
HTS: high-throughput screening;
mAbs: antibodies;
MST: microscale thermophoresis;
SD: standard deviation;
SPR: surface plasmon resonance;
TAMs: tumor-associated macrophages;
TRIC: temperature-related intensity change;

## Notes

### Competing Interest Statement

The authors have declared no competing interest.

