## Supporting Information for "Temperature-Related Intensity Change (TRIC)-based High Throughput Screening Platform for the Discovery of CHI3L1-Targeted Small Molecules"

| **Contents** |  |
| --- | --- |
| Control study for the TRIC-based assay for CHI3L1 binding | S2 |
| MST-based orthogonal validation of potential CHI3L1 hits | S3 |

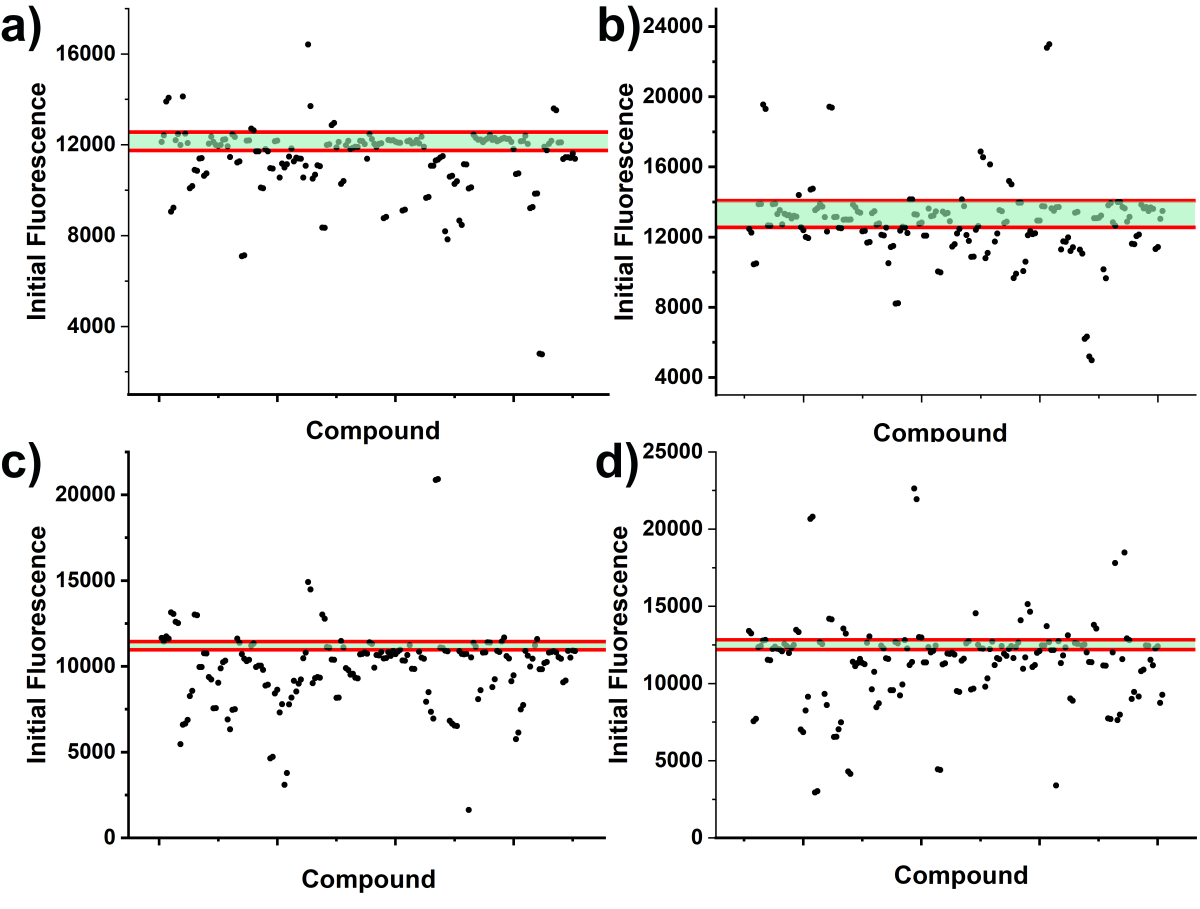

**Figure S1. Control study for the TRIC-based assay for CHI3L1 binding.** Results of the control study at 100 μM. Green region between the red lines indicates the range within 3 × SD from the mean F_norm_ value of negative controls. The graphs show the results from a single experiment.

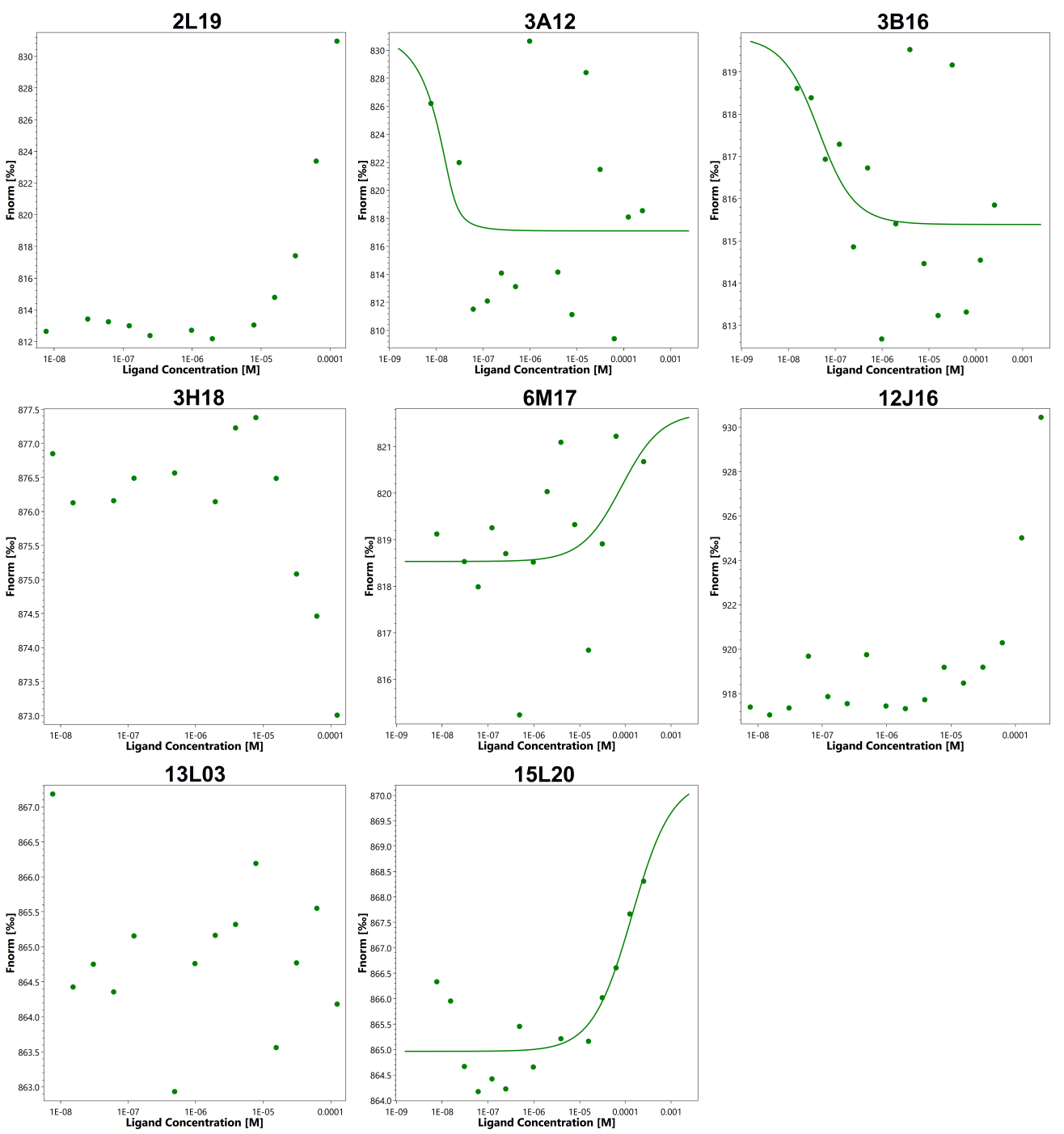

**Figure S2. MST-based orthogonal validation of potential CHI3L1 hits.** Saturation binding curves of hits using MST for the binding of small molecules to CHI3L1.
